# Reduced phosphatases and hyperphosphorylated glutamatergic receptors persist from early to late stages of prion disease

**DOI:** 10.64898/2026.10.14.751571

**Authors:** Alexander J. Rajic, Daniel Ojeda-Juarez, Dipannita Sarkar, Paige Sumowski, Samantha Flores, Mahsa Pourhamzeh, Katrin Soldau, Donald P. Pizzo, Christina J. Sigurdson

**Author notes:** Corresponding author: Christina J. Sigurdson Department of Pathology UC San Diego, 9500 Gilman Dr., La Jolla, CA 92093, USA. Equal contribution (Alexander J. Rajic, Daniel Ojeda-Juarez, Dipannita Sarkar).

## Abstract

Synapse loss occurs early in prion disease, yet the intracellular signaling driving this dysregulation remains poorly understood. Evidence of neuronal hyperactivity was reported in humans with sporadic and familial CJD as well as in the early stages of prion disease in mice (strain 22L). Here we perform a time course study of prion-infected mice (strain RML) and show an increase in the immediate-early gene, *Arc*, suggesting this is a conserved response to prions. We also observed a persistent decrease in phosphatase abundance (PP1α and PP2B) in the cerebral cortex starting at the earliest timepoint (preclinical). At this stage, mGluR5 dimers and phosphorylated AMPA receptors were increased. By clinical disease onset, mGluR5 dimers declined and activity shifted toward NMDA receptor activation, marked by GluN2B hyperphosphorylation. Together, these findings suggest that early phosphatase reduction accompanies neuronal hyperactivity, and precedes substantial inflammation. Elucidating the mechanisms driving this early phosphatase loss and its role in neuronal hyperactivity may reveal novel therapeutic targets for prion disease.

**Significance statement:** Synapse loss and neuronal hyperactivity are early features of prion and Alzheimer’s disease (AD), yet the underlying molecular drivers remain poorly understood. In the current time course study, we reveal elevated levels of neuronal activity protein, Arc, at early, mid, and late timepoints of prion disease. Concurrently, we reveal a persistent and early reduction in key phosphatases (PP1α and PP2B). Additionally, we observed hyperphosphorylation of glutamatergic receptors starting from subclinical stages, before overt neuroinflammation or neuronal loss occur. Furthermore, we uncover a disease-stage-dependent transition from early AMPAR/mGluR5 alterations to late-stage NMDAR involvement. Understanding the mechanisms driving early phosphatase loss and neuronal hyperactivity could reveal novel therapeutic targets for prion disease.

## Introduction

Synapse loss is an early feature of prion-induced neurodegeneration (Belichenko et al., 2000; Jeffrey et al., 2000; Brown et al., 2001; Ferrer, 2002; Cunningham et al., 2003; Gray et al., 2009). Prion-affected patients show aberrant neuronal activity, including increased resting-state connectivity on fMRI (Paoletti et al., 2022), periodic sharp wave complexes on EEG (Wieser et al., 2006), and myoclonus (Sequeira et al., 2022). Additionally, the neuronal activity protein, Arc, is increased in CJD-affected brain (Ojeda-Juarez et al.). Arc is also increased in prion-infected mice starting from preclinical disease stages, and pronounced long, concave post-synapses further support heightened neuronal activity (Siskova et al., 2013; Ojeda-Juarez et al., 2022; Lawrence et al., 2023). Thus, a possible mechanism driving synapse loss is persistent neuronal hyperactivity, yet the alterations in glutamatergic receptors and intracellular signaling that may drive hyperactivity are poorly understood.

The homeostatic control of structural plasticity in the brain is governed in part by the opposing activities of protein kinases (PKs) and protein phosphatases (PPs) (Woolfrey and Dell’Acqua, 2015; Khan et al., 2021). This kinase-phosphatase balance acts as a molecular switch for activity-dependent changes, where PKs typically drive positive modulations such as long-term potentiation (LTP), whereas PPs facilitate negative modulations such as long-term depression (LTD) (Zhabotinsky, 2000; Woolfrey and Dell’Acqua, 2015; Khan et al., 2021). Key mediators of synaptic signaling include Ca^2+^-dependent enzymes such as protein kinase C (PKC) and Ca^2+^/calmodulin-dependent protein kinase II (CaMKII), which are recruited to the postsynaptic density following Ca^2+^ influx through N-methyl-D-aspartate receptors (NMDARs) (Lisman et al., 2012; Wu et al., 2025). These kinases regulate the phosphorylation of synaptic substrates and scaffold proteins to modify synaptic strength and morphology (Colbran and Brown, 2004; Wu et al., 2025). Conversely, protein phosphatases, including protein phosphatase 1 (PP1), protein phosphatase 2A (PP2A), and the Ca^2+^-activated phosphatase, PP2B (calcineurin), provide a counter-regulation by dephosphorylating synaptic substrates, thereby maintaining neuronal signaling within a functional range (Mulkey et al., 1994; Hassan et al., 2024).

This kinase – phosphatase enzyme balance may become perturbed in neurodegenerative disease, transitioning neurons from physiological activity to hyperactivity and eventual synapse loss (Gong et al., 2006). In Alzheimer’s disease (AD), a shift toward aberrant phosphorylation is driven by the overactivation of kinases such as glycogen synthase kinase (GSK)-3β, cyclin-dependent kinase 5 (CDK5), and PKC, and a concurrent reduction in the activity of major phosphatases, including PP1 and PP2A (Gong et al., 2000; Gong et al., 2006). Specifically, PP2A is responsible for the majority of brain Ser/Thr phosphatase activity and serves as the primary tau phosphatase; its downregulation in AD has been shown to facilitate the hyperphosphorylation of tau at sites such as Thr231 and Ser262, impairing microtubule stability (Gong et al., 2000; Liu et al., 2005; Sontag and Sontag, 2014). Emerging evidence suggests that misfolded protein aggregates in other proteinopathies similarly trigger aberrant signaling cascades that disrupt kinase-phosphatase homeostasis (Soto, 2003; Wu et al., 2025).

Prion disease is characterized by the conformational conversion of the cellular prion protein (PrP^C^) into a pathological, β-sheet-rich aggregate, PrP^Sc^ (Kraus et al., 2021; Li et al., 2022). As a glycosylphosphatidylinositol (GPI)-anchored protein, PrP^C^ lacks an intracellular domain and requires transmembrane partners to initiate signal transduction. PrP^C^ has been shown to interact with a diverse array of receptors, including epidermal growth factor receptor (EGFR) (Llorens et al., 2013), G protein-coupled receptors (GPCRs) (Haas et al., 2016), low-density lipoprotein receptor-related protein-1 (LRP1) (Mantuano et al., 2020), neural cell adhesion molecule 1 (NCAM1) (Schmitt-Ulms et al., 2001), and NMDARs (Khosravani et al., 2008). In AD models, the interaction between PrP^C^, Aβ oligomers (Aβο), and mGluR5 triggers a Ca^2+^ response that activates Fyn kinase, leading to the phosphorylation of NMDAR subunit, GluN2B, at Y1472 (Um et al., 2012; Um et al., 2013). Similarly, PrP^Sc^ triggers PrP^C^-dependent signaling, including

NMDAR-mediated Ca^2+^ influx and the subsequent activation of p38 MAPK (Fang et al., 2018), as well as PKC (Lin et al., 2025). However, the temporal changes in PK and PP activity as prion disease progresses, and how this enzyme balance correlates with established markers of neuronal hyperactivity, including Arc, remain to be elucidated. Here we performed a time course study of prion-infected mice (strain RML) and discovered a persistent reduction in phosphatases in the cerebral cortex starting from early disease, when Arc and phosphorylated AMPA receptors were increased.

## Results

### Early prion disease (40%): Mild astro-and microgliosis and decreased synaptophysin

To identify early prion-induced synaptic signaling alterations prior to severe astro- and microgliosis or synapse loss, we performed a time course prion infection study. Wild-type (WT) (C57BL/6) mice were inoculated intracerebrally with RML-infected or uninfected (mock) brain homogenate, and brains were collected at three time points (tp), approximately: (i) 40% [64 days post-inoculation (dpi)], (ii) 70% (130 dpi), and (iii) 100% (179 dpi), corresponding to subclinical, clinical, and terminal disease, respectively. The cerebral cortex is prominently affected during RML infection (Carroll et al., 2016). At the 40% tp, scant PrP^Sc^ was evident (Fig.1, Extended data Fig.1-1), there was little to no vacuolation, and astrocytic and microglial proteins (GFAP and Iba1, respectively) were subtly increased in the cerebral cortex by immunohistochemistry (IHC) and western blot (Fig. 1). Furthermore, neurons, dendrites, and synapses, assessed by NeuN, MAP2, and synaptophysin immunolabeling, respectively, were unaltered (Extended data Fig. 1-2). By the 70% tp, PrP^Sc^ was diffusely distributed throughout the cortex, with moderate astro- and microgliosis. By western blot, GFAP and Iba1 were also markedly elevated (Fig.1) and were further increased in the terminal brain (Fig. 1).

**Figure 1.**
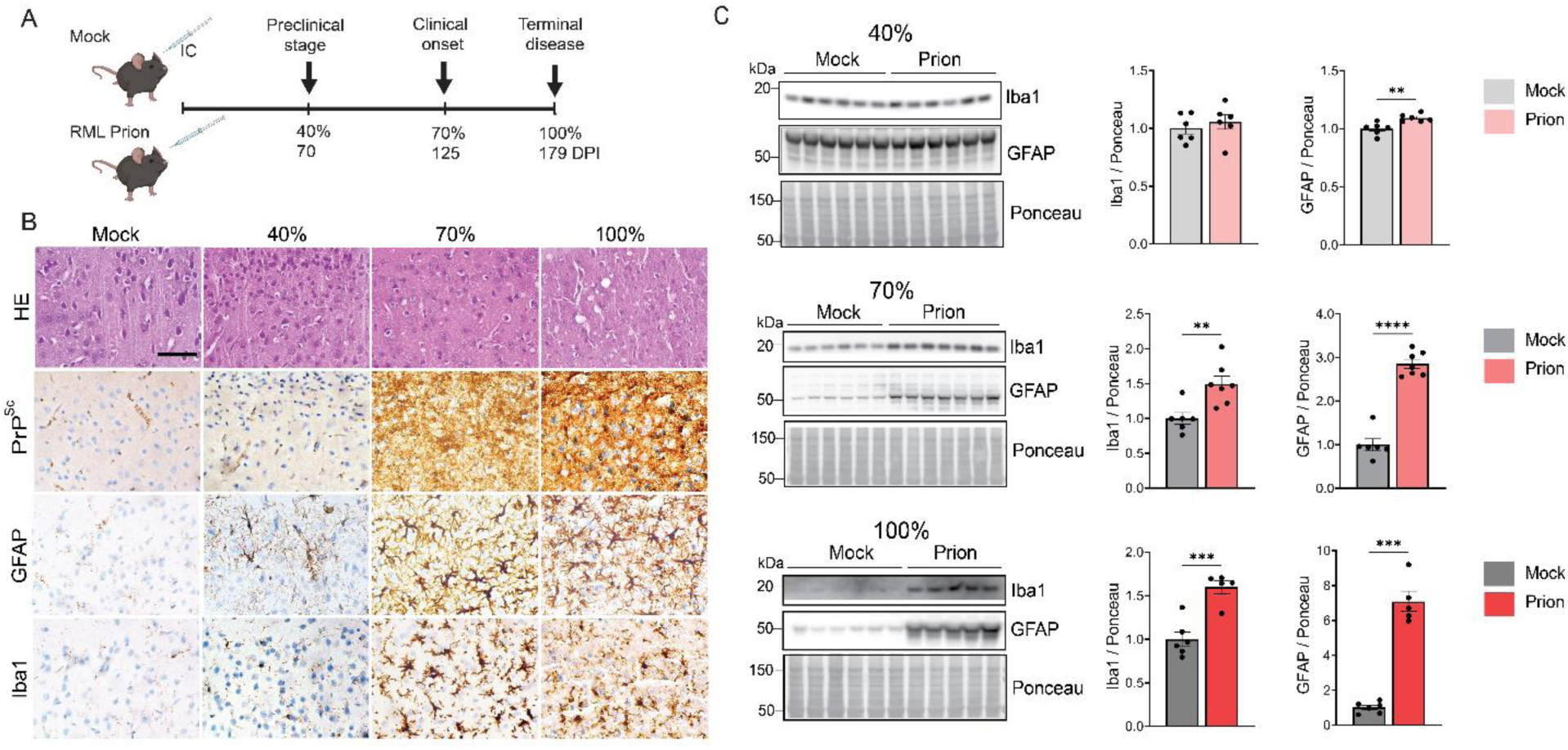
Prion-infected mice show mild astrogliosis prior to PrP^Sc^ accumulation or neuronal spongiform degeneration. (A) Schematic of time course experiment showing WT mice inoculated with uninfected or prion-infected brain homogenate (RML strain). (B) Representative images of cortex (layers II-III) from uninfected or prion-infected WT mice stained with hematoxylin and eosin (HE), or immunolabeled for PrP^Sc^, GFAP, or Iba1. Three time points were analyzed (40%, 70%, and 100%). (C) Immunoblot and quantification show GFAP and Iba1 at 40%, 70%, and 100% time points. 40%: N = 6 per group, 70%: N = 6 (mock) and 7 (prion), 100%: N = 6 (mock) and 5 (prion). Mean ± SEM. Unpaired, 2-tailed *t* test with Welch’s correction. \*\**P*<0.01, \*\*\**P*<0.001, \*\*\*\**P*<0.0001.

To more accurately quantify synaptic proteins, we measured the pre- and post-synaptic proteins, synaptophysin and PSD95, respectively, by western blot. At the 40% tp, synaptophysin was reduced (0.7-fold) while PSD95 was unchanged (Fig. 2). The cytosolic pre-synaptic protein, synapsin-1, was increased (1.4-fold). By the 70% tp, synaptophysin and synapsin-1 were similar to the controls, however by terminal disease, both were markedly reduced (0.8-and 0.7-fold, respectively) (Fig. 2). PSD95 was reduced at the 70% and 100% tp (0.7- and 0.9-fold, respectively) (Fig. 2). Because the 40% tp lacked overt inflammatory changes, neuronal loss, or synaptic loss, we selected this subclinical window to investigate early alterations in immediate early genes (IEGs) as indicators of synaptic activity in the cerebral cortex.

**Figure 2.**
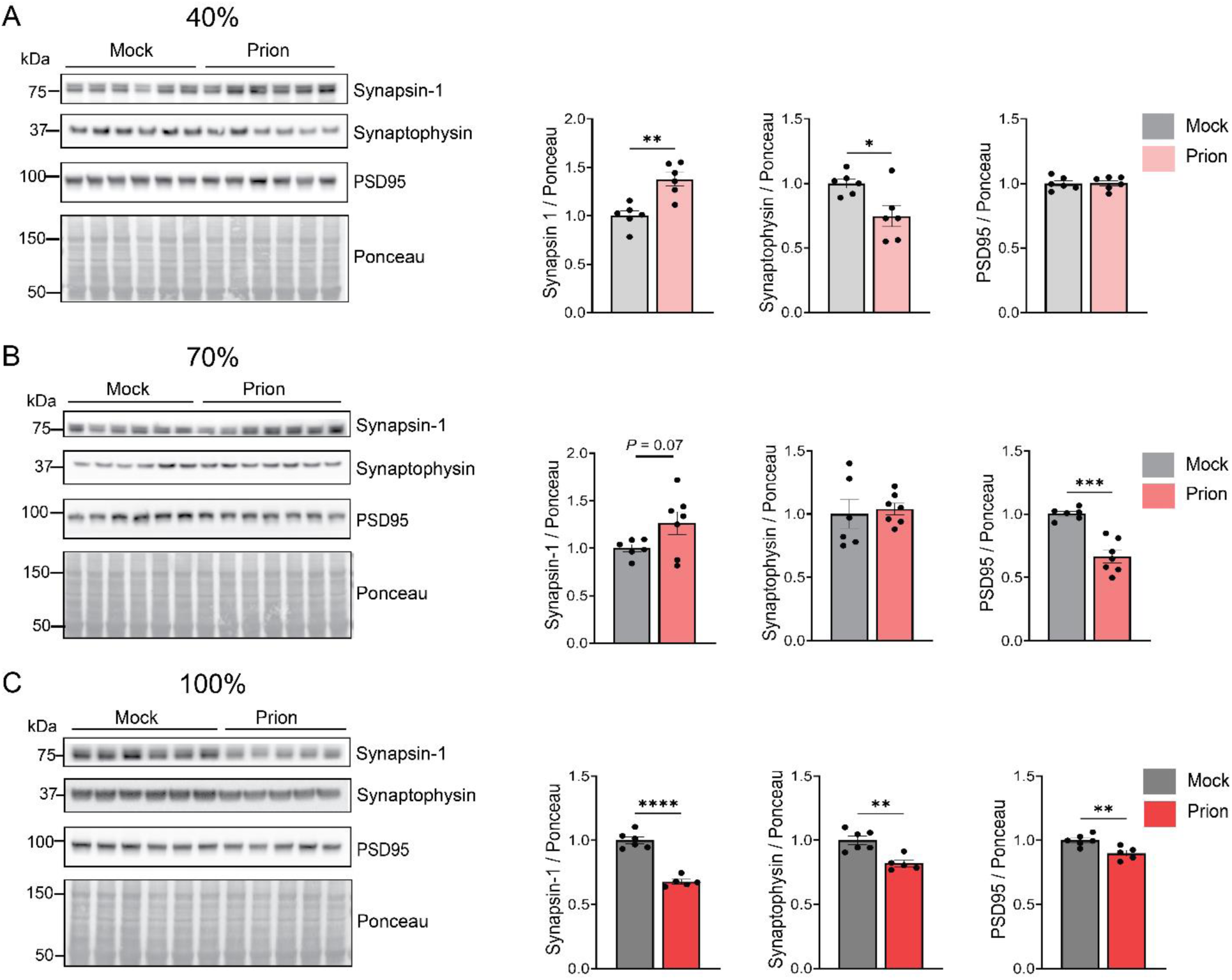
Pre-synaptic proteins are altered in the prion-infected cerebral cortex at the 40% timepoint, while pre-and post-synaptic proteins are reduced by terminal disease. (A) Immunoblots and quantification of mock and prion-infected mouse cortex show pre-synaptic cytoplasmic and vesicle proteins, synapsin-1 and synaptophysin, respectively, and post-synaptic protein, PSD95. 40%: N = 6 per group, 70%: N = 6 (mock) and 7 (prion), 100%: N = 6 (mock) and 5 (prion). Mean ± SEM. Unpaired, 2-tailed *t* test with Welch’s correction. \**P*<0.05, \*\**P*<0.01, \*\*\**P*<0.001, \*\*\*\**P*<0.0001.

### Arc increases while Egr1 decreases in early prion disease

IEGs are expressed in response to increased neuronal activity, and Arc was previously observed to be increased in early disease in mice infected with 22L prions (Ojeda-Juarez et al., 2022). To determine whether IEGs were altered consistent with heightened neuronal activity, we measured Arc, Egr1, c-Fos, and Homer1 by western blot. There was variability at the 40% tp, as Arc was modestly elevated (1.1-fold), while Egr1 was reduced (0.7-fold) (Fig. 3).

**Figure 3.**
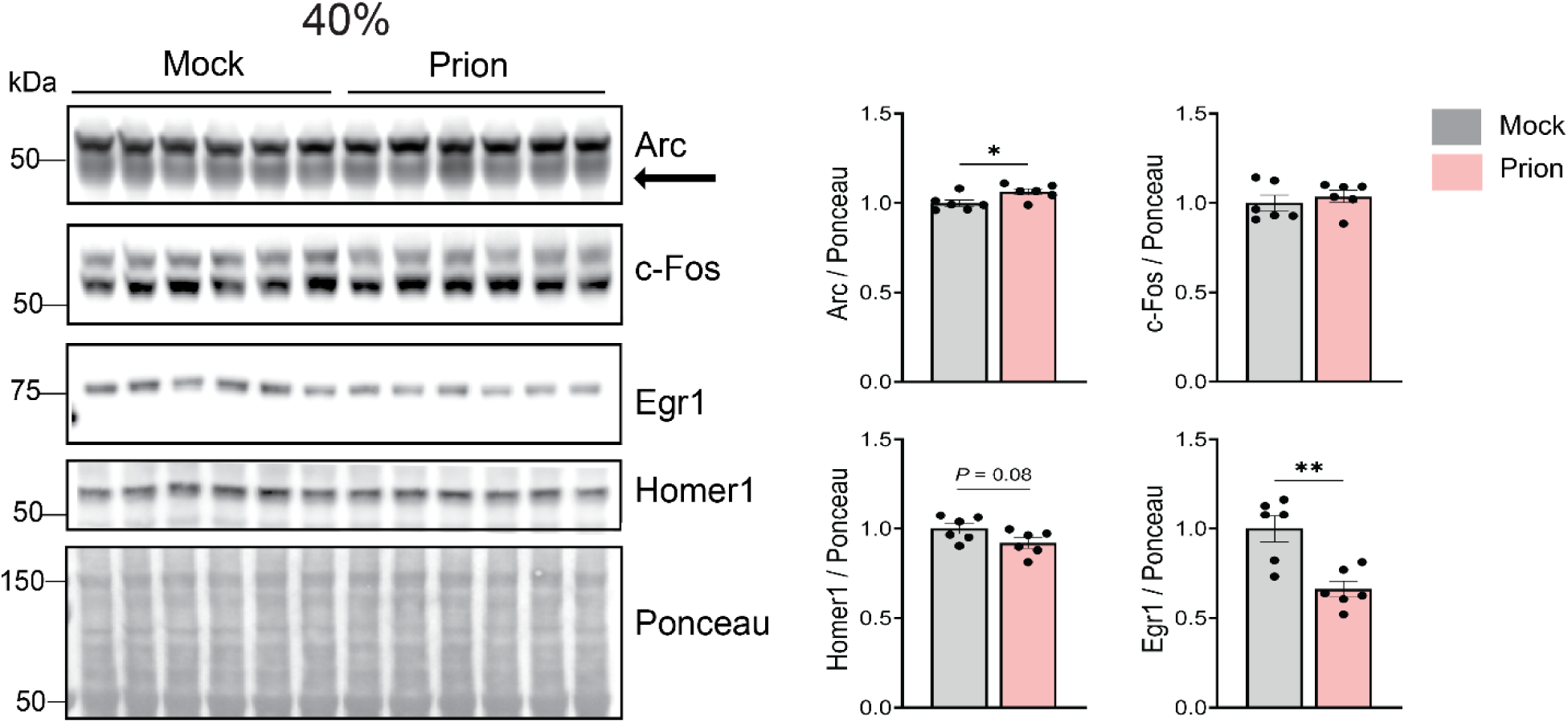
IEG proteins, Arc and Egr1, are altered in the prion-infected cerebral cortex at the 40% timepoint. Immunoblots and quantification of four IEGs: Arc, c-Fos, Homer1, and Egr1. N = 6 per group. Mean ± SEM. Unpaired, 2-tailed *t* test with Welch’s correction. \**P* < 0.05 and \*\**P* < 0.01.

### An early increase in synaptic surface receptors: mGluR5 dimers and phosphorylated AMPA receptors

Arc may be increased as a result of increased synaptic activity following NMDA receptor stimulation (Chen et al., 2017; Yang et al., 2023) or by mGluR5-mediated activity (You et al., 2012; Um et al., 2013; Hu et al., 2014; Chen et al., 2024; Lin et al., 2025). At the 40% tp, mGluR5 dimers were increased (1.1-fold), indicating elevated surface expression (Fig. 4), which may underlie the increase in Arc. In contrast, GluN2B subunits phosphorylated at S1303 (pGluN2B-S1303) were decreased (0.8-fold) (Fig. 4), consistent with decreased Ca^2+^ permeability (Liao et al., 2001) and suggestive of reduced NMDAR channel conductance at an early timepoint.

**Figure 4.**
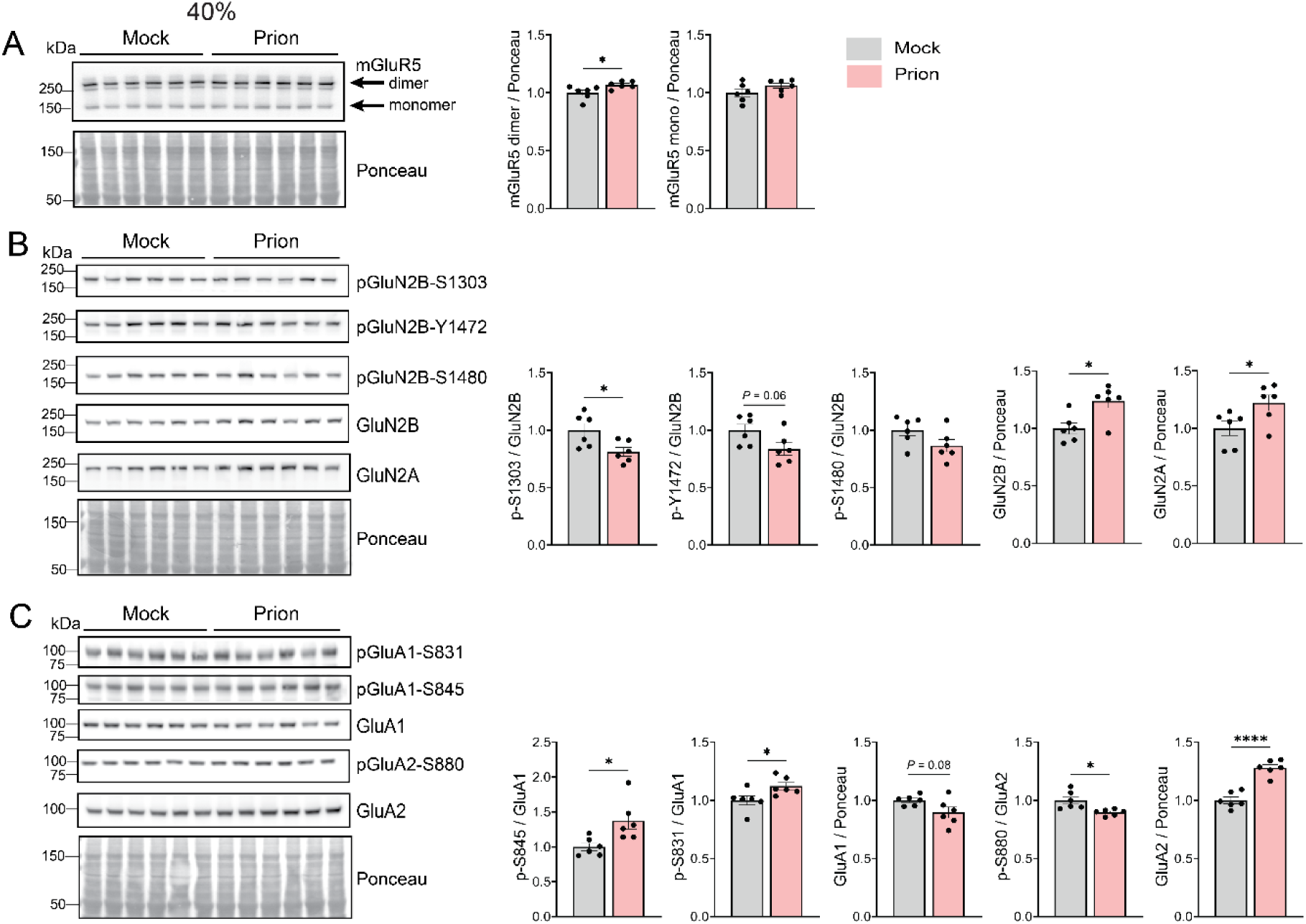
Phosphorylated glutamatergic receptor levels are altered in prion-infected cerebral cortex at the 40% timepoint. Immunoblots and quantification of (A) mGluR5 monomers and dimers, (B) NMDA receptors, GluN2A, and GluN2B, and key NMDAR phosphosites (GluN2B: S1303, Y1472, and S1480). (C) AMPA receptor subunits, GluA1 and GluA2, and key phosphosites (GluA1: S845, and S831; GluA2: S880). N = 6 per group. Mean ± SEM. Unpaired, 2-tailed *t* test with Welch’s correction. \**P*<0.05 and \*\*\*\**P*<0.0001.

Arc contributes to homeostatic synaptic downscaling in less active synapses by modulating GluA1 endocytosis (Chowdhury et al., 2006; Shepherd et al., 2006; Bramham et al., 2008). Given the increase in Arc, we measured the phosphorylation status of GluA1 and GluA2 receptors. At the 40% tp, phosphorylated GluA1 at S831 and S845 were increased by 1.1- and 1.4-fold, respectively (Fig. 4), suggestive of increased single channel conductance (Summers et al., 2019), and AMPAR primed for synaptic membrane insertion (Esteban et al., 2003; He et al., 2009). Phosphorylated GluA2 at S880 was decreased (0.90-fold), while total GluA2 levels were increased (1.3-fold) (Fig. 4), suggestive of increased synaptic membrane localization of GluA2 (Matsuda et al., 1999), although surface localization was not directly assessed. Together, these changes in phosphorylated glutamate receptors are consistent with a shift toward AMPAR states that favor membrane retention of both Ca^2+^-permeable (GluA1) and impermeable (GluA2) AMPAR subunits, suggestive of upscaling and strengthening synapses, tempered by limiting Ca^2+^ influx.

### Reduced phosphatase levels (PP2B and PP1a) at the 40% tp

To test the upstream signaling that leads to alterations in AMPAR and NMDAR subunit phosphorylation, we next measured relevant kinases (PKA, PKC, CaMKII, and Fyn) and phosphatases (PP1α, PP2A, PP2B, STEP61) by western blot. Protein kinase A (PKA) phosphorylates GluA1 at S845 (Mammen et al., 1997; Dell’anno et al., 2013) and PP2B dephosphorylates this site (Sanderson et al., 2012). PKC (Roche et al., 1996; Jenkins and Traynelis, 2012) and CaMKII (Barria et al., 1997; Mammen et al., 1997) phosphorylate GluA1 at S831, GluA2 at S880, and GluN2B at S1303. PP1α dephosphorylates GluA1 at S831 and at S845 (Huang et al., 2001; Hu et al., 2007), and GluN2B at S1303 (Huang et al., 2001; Hu et al., 2007; Raveendran et al., 2009; Mukherjee et al., 2010b; Farinelli et al., 2012), whereas PP2A dephosphorylates GluA2-S880 (Launey et al., 2004), and GluN2B-S1303 (Raveendran et al., 2009).Finally, Fyn kinase phosphorylates GluN2B at Y1472 (Nakazawa et al., 2001), while both Fyn and GluN2B-1472 are dephosphorylated by STEP61 (Nguyen et al., 2002; Goebel-Goody et al., 2009).

At the 40% tp, there were no changes in phospho-PKA substrates or phospho-PKC substrates, or in pCaMKII-T286 (Extended data Fig. 5-1). However, PP2B was markedly decreased (0.7-fold) and PP1α was modestly decreased (0.9-fold) (Fig. 5). These findings raise the possibility that the increased AMPAR phosphorylation may reflect reduced phosphatase level and activity rather than an increase in kinase activity, although phosphatase activity was not directly measured.

**Figure 5.**
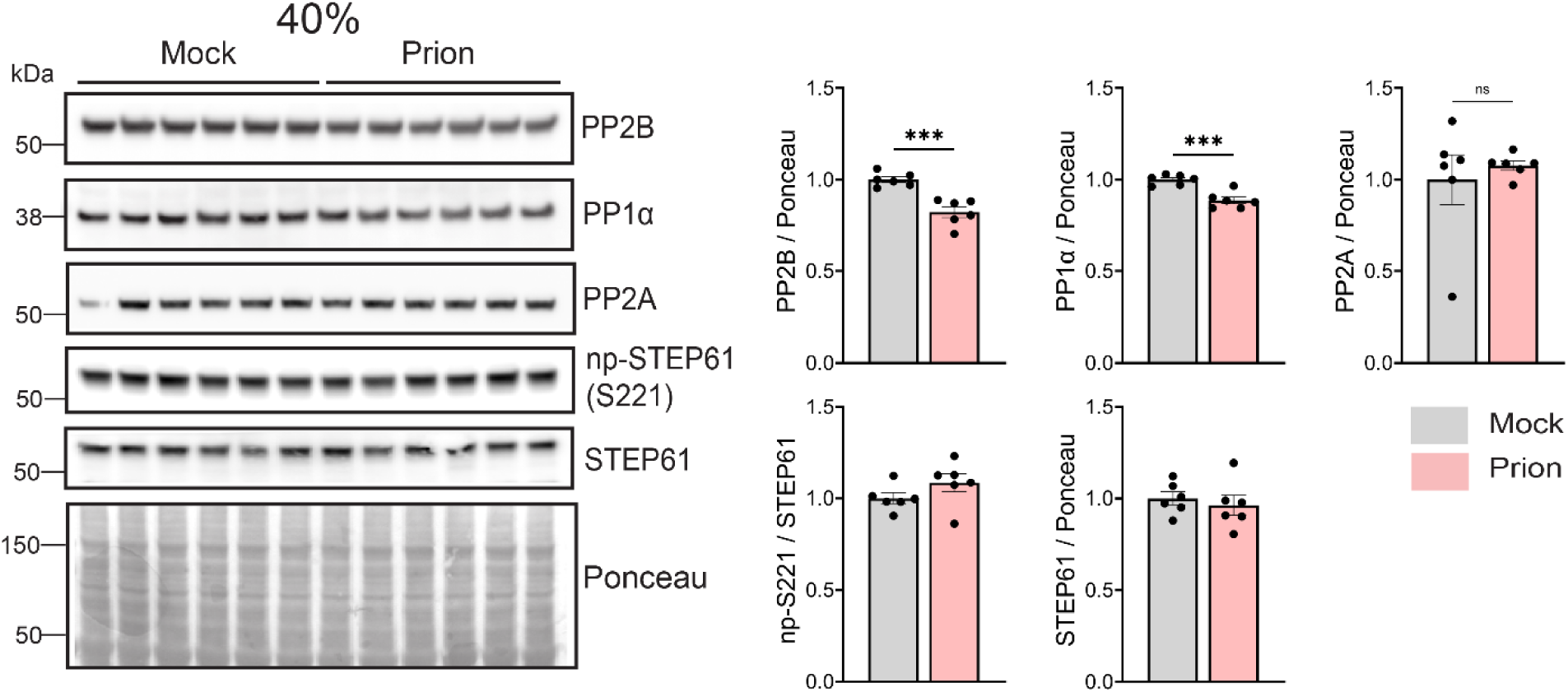
Phosphatases, PP2B and PP1α, are reduced in the cerebral cortex at the 40% timepoint. (A) Immunoblots and quantification of serine/threonine protein phosphatases, PP2B, PP1α, PP2A, and STEP61. N = 6 per group. Mean ± SEM. Unpaired, 2-tailed *t* test with Welch’s correction. \*\*\**P*<0.001.

### Phosphatase levels remain reduced, while PKA substrates and glutamate receptor phosphorylation increases, in clinical stages of disease

We next evaluated neuronal activity proteins, glutamatergic receptors, and kinases and phosphatases at clinical onset (70% tp) and at terminal disease (100% tp). By the 70% tp, mice developed early clinical signs such as kyphosis and stiff tails. At both the 70% and 100% tp, Arc was elevated (1.1- and 1.2-fold, respectively) (Extended data Fig. 4-1), which may reflect sustained neuronal hyperactivity or decreased degradation, as mGluR5 dimers were reduced (0.9-fold at 70%; 0.4-fold at 100%) (Extended data Fig. 4-1). Concurrently, pGluN2B-Y1472 was markedly increased (1.4-fold) by the 70% tp, but was transient, returning to baseline by the 100% tp (Extended data Fig. 4-1). By the 100% tp, pGluA1-S845 was increased by 2.2-fold (Extended data Fig. 4-1). Together, these data are suggestive of elevated glutamatergic activity in the cerebral cortex during clinical prion disease.

We next measured the kinase and phosphatase levels at the later time points. By the 70% tp, Fyn kinase was increased (1.3-fold) (Extended data Fig. 6-1), consistent with the elevated phosphorylation of its substrate, GluN2B at Y1472 (Extended data Fig. 4-1). By the 100% tp, phospho-PKA substrates were increased (2.9-fold) (Extended data Fig. 6-1), consistent with the increased phosphorylation of PKA substrate, GluA1-S845.

In contrast, reduced phosphatase levels persisted through later disease, but with a shifting pattern. By the 70% tp, PP1α and PP2A were markedly decreased (0.5- and 0.3-fold, respectively) (Fig. 6). By the 100% tp, reductions spanned three of the four phosphatases, with PP1α, PP2B, and STEP61 reduced by approximately 30% (Fig. 6). Together, these data are suggestive of a sustained reduction in phosphatase levels from early to terminal stages of disease, with increased PKA substrate phosphorylation at terminal disease, which together could favor maintaining glutamatergic receptors in a hyperphosphorylated state.

**Figure 6.**
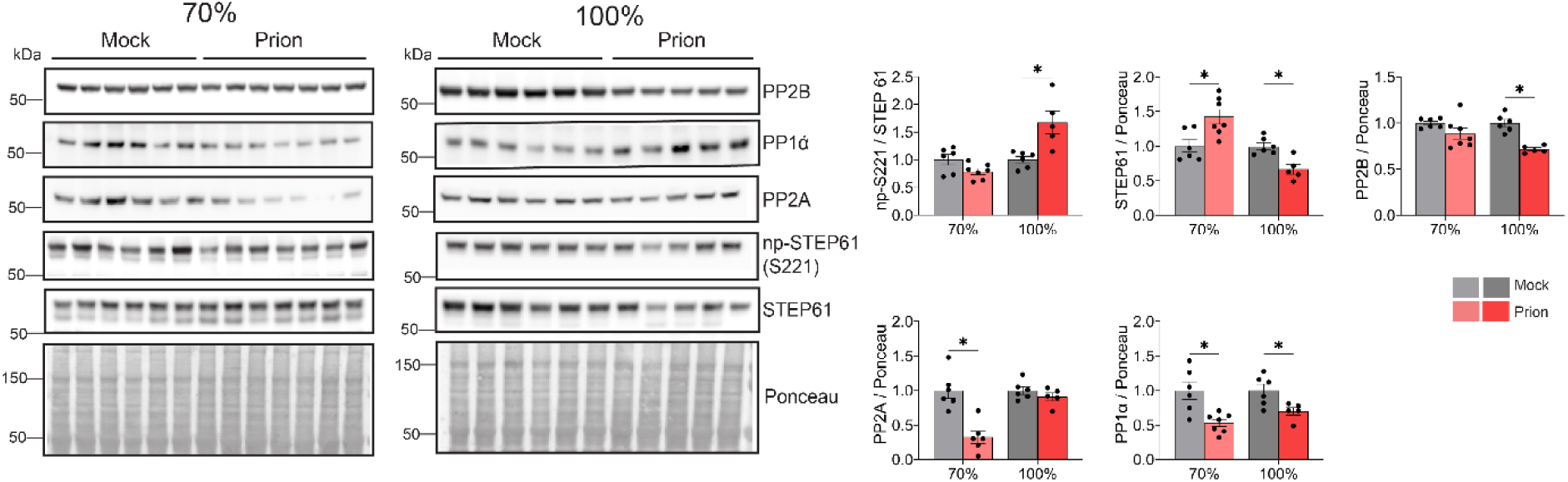
Phosphatases are reduced in the cerebral cortex at clinical stages of prion disease (70% and 100%). (A) Immunoblots of phosphatases at 70% and 100% (left) and quantification at all time points (right). 70%: N = 6 (mock) and N = 7 (prion); 100%: N = 6 (mock) and N = 5 (prion). Mean ± SEM. Unpaired, 2-tailed *t-test* with Welch’s correction. \**P*<0.05.

## Discussion

Glutamate receptor phosphorylation has been previously shown to be altered in prion disease (Bertani et al., 2017; Ojeda-Juarez et al., 2022; Lawrence et al., 2023; Lin et al., 2025), but how the associated kinases and phosphatases change has been unclear. In the present study, we identify an early reduction in protein phosphatases in the cerebral cortex that persisted through three disease stages. Specifically, we showed a marked reduction in the serine/threonine phosphatases, PP2B, PP1α, and PP2A at the earliest timepoint, when Arc levels were also elevated (40%, 64 dpi). These alterations were accompanied by an early increase in phosphorylated AMPARs and mGluR5 dimers, and a later increase in phosphorylated NMDARs, all with the potential to increase cytoplasmic Ca^2+^ (Reiner and Levitz, 2018). The chronic reduction of phosphatases supports a model of kinase-phosphatase imbalance that may facilitate an early drive of synaptic plasticity toward hyperactivity prior to neurodegeneration.

At the preclinical (40%) timepoint, we observed an increase in Arc and phosphorylated AMPAR, accompanied by scant PrP^Sc^ accumulation, minimal to no neuronal vacuolation or synapse loss, and mild astrogliosis in the cerebral cortex of mice infected with the RML prion strain, consistent with increased neuronal activity prior to overt neuropathology. Previous studies have reported increased Arc in the brains of mice infected with the 22L prion strain (Ojeda-Juarez et al., 2022), and in patients affected by genetic and sporadic prion disease (Ojeda-Juarez et al., 2026); here we extend those studies to show the increase in Arc in mice infected with a different prion strain. Congruent with increased neuronal activity, patients in the early-to mid-stages of sCJD show increased functional connectivity in the ventral and dorsal default mode networks by fMRI (Paoletti et al., 2022). Together, these findings suggest that neuronal hyperactivity may be among the earliest disease alterations, occurring following infection with two different mouse prion strains, as well as in both sporadic and familial human prion disease.

While elevated Arc persisted throughout disease, we observed a distinct time-dependent switch in glutamatergic receptor hyperphosphorylation. In early disease, we observed increased AMPAR phosphorylation at GluA1 residues S845 and S831, as well as reduced GluA2 phosphorylated at S880, suggestive of increased synaptic membrane insertion and retention of both Ca^2+^-permeable and impermeable AMPARs and a pronounced AMPAR-mediated excitatory drive. As the disease progressed, the glutamate receptor signaling shifted to include a stronger NMDA receptor-driven excitatory state. At symptom onset (70%), phosphorylation of GluN2B Y1472 increased, suggestive of greater stabilization of synaptic GluN2B and enhanced NMDA receptor signaling coinciding with worsening clinical signs. Consistent with these observations of hyperphosphorylated glutamate receptors, acute toxicity studies have shown that inhibitors of AMPAR and NMDAR reduce spine retraction in hippocampal neurons 24 hours post-PrP^Sc^ exposure (Fang et al., 2018). The observed hyperphosphorylated AMPAR and NMDAR would be predicted to increase Ca^2+^ influx and drive plasticity changes (Makino et al., 2011; Diering and Huganir, 2018) that could contribute to excitotoxicity and neuronal death. The mechanisms regulating this transition from AMPAR to NMDAR activation require further investigation.

We observed an early increase in mGluR5 dimers followed by a steep decline at the pre-symptomatic and terminal stages of disease, congruent with recent reports in RML-infected mice (Wang et al., 2026). This reduction of mGluR5 dimers in late prion disease appears to be strain- and species-independent, as decreased mGluR5 (combined monomers and dimers) occurs across multiple mouse prion strains (22L (Ojeda-Juarez et al., 2022), ME7, 139A (Hu et al., 2022)), as well as in hamsters (Ha-263K) (Hu et al., 2022), non-humans (Wang et al., 2026), and humans (sCJD) (Ojeda-Juarez et al., 2022). Whether this reduction is compensatory to prevent prolonged elevated cytoplasmic Ca^2+^, or whether it is pathologic and advances disease progression remains unclear. Interestingly, pharmacological inhibition of Group 1 metabotropic receptors (mGluR1 and mGluR5) prolonged survival in prion-infected mice by inhibiting both receptors (Goniotaki et al., 2017), yet the contribution and cell-type specific effects of individual receptors remain to be elucidated. Additionally, the mechanisms and consequences of mGluR5 loss in prion disease are unknown and warrant further investigation.

The PP2B, PP1α, and PP2A levels were decreased without evidence of kinase activity changes in early disease. This profile suggests an imbalance between kinases and phosphatases that skews signaling toward phosphorylation-driven receptor trafficking and surface stabilization, which may bias glutamatergic receptor dynamics toward a more excitable state, although whether phosphatase loss equates to reduced activity remains to be elucidated. This finding is particularly important as it stands in paradoxical contrast to reported increases in PP2B activity in human neuroblastoma cell lines treated with the PrP (106-126) peptide (Hong et al., 2020; Moon et al., 2021). Additionally, PP2B activity was increased in prion-infected whole brain, and inhibition of PP2B with inhibitor, FK506, prolonged the survival of RML-infected mice (Mukherjee et al., 2010a), however whether the levels were unaltered is unclear. Understanding the mechanisms driving the observed early phosphatase loss and its role in neuronal hyperactivity could reveal novel therapeutic targets for prion disease.

Our results establish a molecular basis for glutamatergic pathology in prion disease, contrasting early phosphatase alterations in prion-infected brains with kinase-phosphatase imbalance observed in Alzheimer’s disease. Our interpretations of western blot data are constrained by the use of bulk tissue measurements, which do not directly indicate receptor localization, electrophysiological activity, or Ca^2+^ flux. Further studies using enzyme activity assays and cell-type-specific approaches will be necessary to elucidate the mechanisms driving these changes. Nevertheless, the present findings identify stage-dependent glutamatergic alterations as an early and progressive feature of prion disease in the cerebral cortex, along with disrupted phosphatase levels, providing a framework for future studies aimed at defining the mechanisms that drive synaptic dysfunction and neurodegeneration. More broadly, the stage-specific synaptic dysregulation may help clarify signaling pathway-specific vulnerability during disease progression and inform the development of stage-specific interventions.

### Materials and Methods Prion inoculation of mice

C57BL/6J mice (6–8 weeks old) were maintained under specific pathogen-free conditions on a 12:12 h light/dark cycle with ad libitum access to food and water. Mice were anesthetized with ketamine and xylazine and inoculated intracerebrally into the right parietal cortex with 30 µl of 1% (w/v) RML prion-infected brain homogenate or uninfected (mock) brain homogenate. Mice were monitored for clinical signs, including kyphosis, ataxia, and stiff tail. Animals were euthanized at three predetermined timepoints: early disease (64 dpi, approximately 40% tp), clinical onset (130 dpi, approximately 70% incubation), and terminal disease (175-182 dpi). The incubation period for terminal disease was estimated and can vary between cohorts.

### Ethics statement and institutional approval

All animal procedures were performed in accordance with the recommendations in the Guide for the Care and Use of Laboratory Animals of the National Institutes of Health. The protocol was approved by the Institutional Animal Care and Use Committee (IACUC). All efforts were made to minimize animal suffering and to reduce the number of animals used.

### Immunoblotting

Brain tissues (cortex) were homogenized in RIPA buffer (10% w/v) using a Beadbeater™ tissue homogenizer. Samples were lysed on ice for 30 min using 2% N-lauryl sarcosine in RIPA with protease and phosphatase inhibitors (Complete™ and Phos-STOP™, Roche). Lysates were cleared by centrifugation at 2,000 x g for 10 min at 4°C. Protein concentrations were determined using a bicinchoninic acid (BCA) assay. Equivalent amounts of protein (20–40 µg) were reduced with dithiothreitol (100 mM), resolved by SDS-PAGE using Bis-Tris gels, and transferred to nitrocellulose membranes. Membranes were blocked in casein or BSA in TBS with Tween 20 (TBST) for 30 min, and incubated with primary antibodies overnight at 4°C. Following incubation with HRP-conjugated secondary antibodies, signals were developed using enhanced chemiluminescence (ECL Supersignal West Dura or Femto, ThermoFisher Scientific) and captured using the Fuji LAS 4000 imager. Densitometry analysis was performed using MultiGauge software (Fujifilm), with protein levels normalized to total protein. For mGluR5 dimer analysis, non-reducing conditions were employed to preserve dimeric structures.

### Immunohistochemistry

Formalin-fixed, paraffin-embedded (FFPE) brain sections (5 µm) were utilized for histological and IHC analyses. Sections were deparaffinized and rehydrated through a graded ethanol series. Spongiform change was assessed using hematoxylin and eosin (HE) staining. For glial fibrillary acidic protein (GFAP) (DAKO), ionized calcium binding adaptor molecule 1 (Iba1) (Wako), PrP (SAF84; Cayman Chemical), and NeuN (Abcam), immunolabeling was performed on an automated tissue immunostainer Ventana Discovery Ultra system (Ventana Medical Systems, Tucson, AZ, USA). Antigen retrieval was optimized for each epitope to yield the maximal signal-to-noise ratio. Stains with heat-based antigen retrieval used CC1 (Tris-EDTA-based; pH 8.6) for 40 min at 95° C (NeuN, Iba), while GFAP used protease for 16 min (Protease 2; Ventana). For PrP^SC^, slides were incubated in protease 2 for 20 min, followed by antigen retrieval in CC1 (Tris-based; pH 8.5; Ventana) for 64 min at 95° C. The primary antibodies were incubated on the sections for 32 min at 37° C. Primary antibodies were detected using the OmniMap systems (Ventana; goat anti-rabbit or goat anti-mouse). Antibody presence was visualized using DAB as a chromogen followed by hematoxylin as a counterstain. Slides were rinsed, dehydrated through alcohol and xylene, and cover-slipped.

To determine neuronal density in prion-infected mice at the 40% tp, NeuN-stained cerebral cortices (layers II-III) were imaged using the Olympus EX41 microscope with DP Controller. All images were acquired on the same day using identical spatial calibration. Image acquisition and quantification were performed blinded to experimental group. The number of neurons over area (neuronal density) was determined by manually counting NeuN^+^ in ImageJ using the Cell Counter plugin for three regions of interest of cortex per mouse. The average neuronal density (number of NeuN^+^ cells/ pixel^2^) per mouse was used for statistical analysis.

### Immunofluorescence staining

For synaptic and dendritic analysis, FFPE brain sections were dual-labeled for synaptophysin and MAP2, respectively. Sections were deparaffinized, heated in citrate buffer for epitope retrieval (Sigma) in a pressure cooker for 30 min, quenched in 3% H_2_O_2_ in methanol for 15 min, and blocked in 1X PBS with 0.1% Tween 20 and 5% goat serum. Slides were then incubated with the primary antibodies, synaptophysin (Cell Signaling Technology #5461) and MAP2 (Invitrogen #PA1-10005), overnight at 4 °C. MAP2 was visualized using biotin-conjugated anti-chicken IgG (Jackson ImmunoResearch #703-065-15) (30 min), then streptavidin-488 (Jackson ImmunoResearch #016-540-084) (30 min), while synaptophysin was visualized using anti-rabbit Cy3 (Jackson ImmunoResearch #111-065-003). All slides were mounted using ProLong Gold (ThermoFisher Scientific).

For MAP2 and synaptophysin image quantification from immunolabeled cerebral cortices (layers II-III) full thickness images (Z-stacks) were acquired using the Nikon Eclipse Ti2-E microscope using the laser scanning confocal mode (A1R HD) and a Nikon Plan Apo ʎ 60x NA 1.40 oil objective (3 fields per mouse). Images were deconvolved and Best Focus function was used to select the optimal 2D analysis plane using the NIS elements software (version 5.42.02: High Content Analysis package). ImageJ software was utilized to determine percent of the neuropil area containing dendrites (MAP2) or pre-synapses (synaptophysin). The MAP2 or synaptophysin percent area from each of the three fields was averaged, and the average value from each mouse was used for the statistical analysis.

### Primary antibodies

The following primary antibodies were used: PrP (POM19), GFAP (1:2000, Dako #Z0334), Iba1 (1:1000, Cell Signaling Technology #17198), synaptophysin (1:5000, Cell Signaling Technology #36406), Synapsin1 (1:5000, Cell Signaling Technology #5297), PSD95 (1:5000, Cell Signaling Technology #3450), Arc (1:2000, Proteintech #16290–1-AP), Egr1 (1:1000, Cell Signaling Technology #4154), c-Fos (1:1000, Cell Signaling Technology #2250), Homer1 (1:1000, Cell Signaling Technology #8231), mGluR5 (1:5000, Cell Signaling Technology #55920), GluN2A (1:1000, Cell Signaling Technology #4205), GluN1 (1:500, Cell Signaling Technology #5704), GluN2B (1:1000, Cell Signaling Technology #14544), p-GluN2B Y1472 (1:1000, Cell Signaling Technology #4208), p-GluN2B S1303 (1:1000, Cell Signaling Technology #71335), p-GluN2B S1480 (1:1000, PhosphoSolutions #p1516-1480), anti-phosphorylated GluA1-S845 (1:1000, Cell Signaling Technology #8084); anti-GluA1 (1:1000, Cell Signaling Technology #13185), anti-phosphorylated GluA1-S831 (1:1000, Cell Signaling Technology #75574), p-GluA2 S880 (1:1000, PhosphoSolutions # p1170-880), GluA2 (1:1000, Cell Signaling Technology #13607) PP2B (calcineurin A) (1:1000, Cell Signaling Technology #2614), PP1α (1:1000, Cell Signaling Technology #2582), PP2A (1:1000, Cell Signaling Technology #2041), Fyn (1:1000, Cell Signaling Technology #4023), STEP (1:1000, Cell Signaling Technology #4396), and non-phospho STEP (Ser221)(1:1000, Cell Signaling Technology #5659).

## Statistical analysis

Data are presented as mean ± SEM. Statistical significance was determined using Welch’s t-test for comparisons between two groups or one-way ANOVA followed by Tukey’s HSD post-hoc test for multiple comparisons. No samples were excluded from analysis. *P*-values < 0.05 were considered significant. All statistical analyses were performed using GraphPad Prism.

## Extended data

**Figure 1-1.**
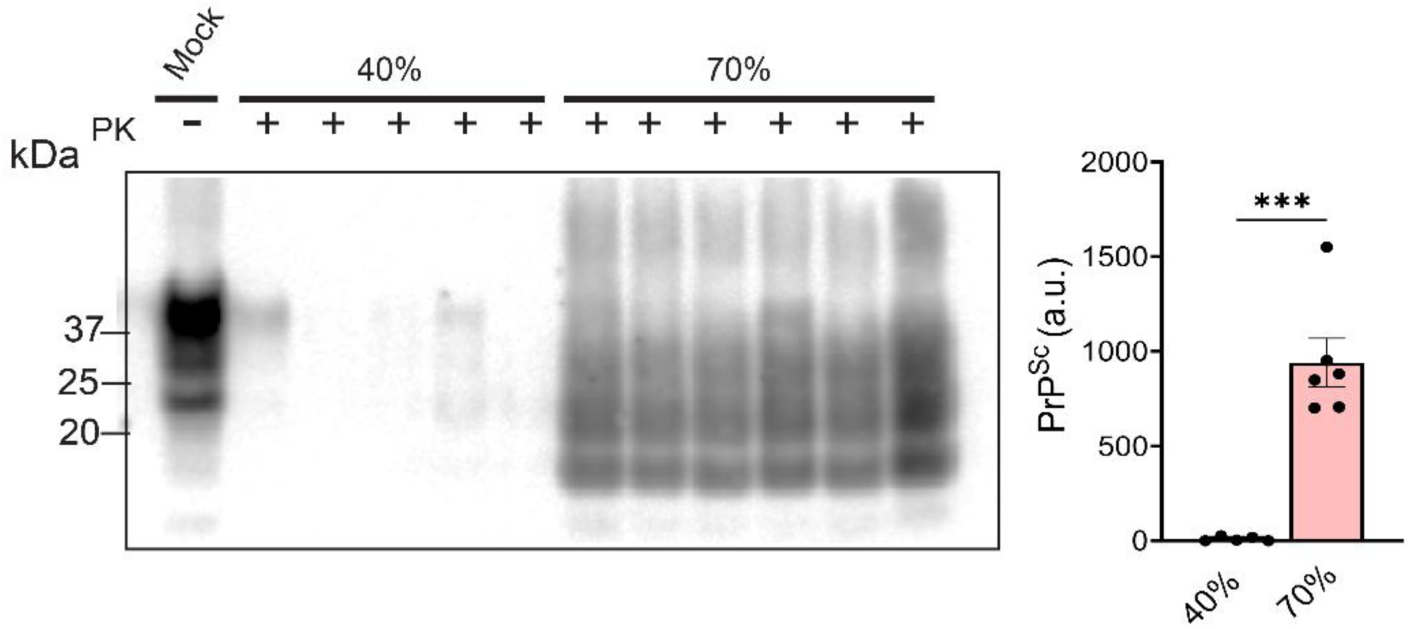
PrP^Sc^ accumulates in the cerebral cortex at the clinical stage of disease (70%). Western blot showing PK-resistant PrP^Sc^ in cortex at the 40% and 70% timepoints. 40%: N = 5, 70%: N = 6. Mean ± SEM. Unpaired, 2-tailed *t* test with Welch’s correction. \*\*\**P*<0.001.

**Figure 1-2.**
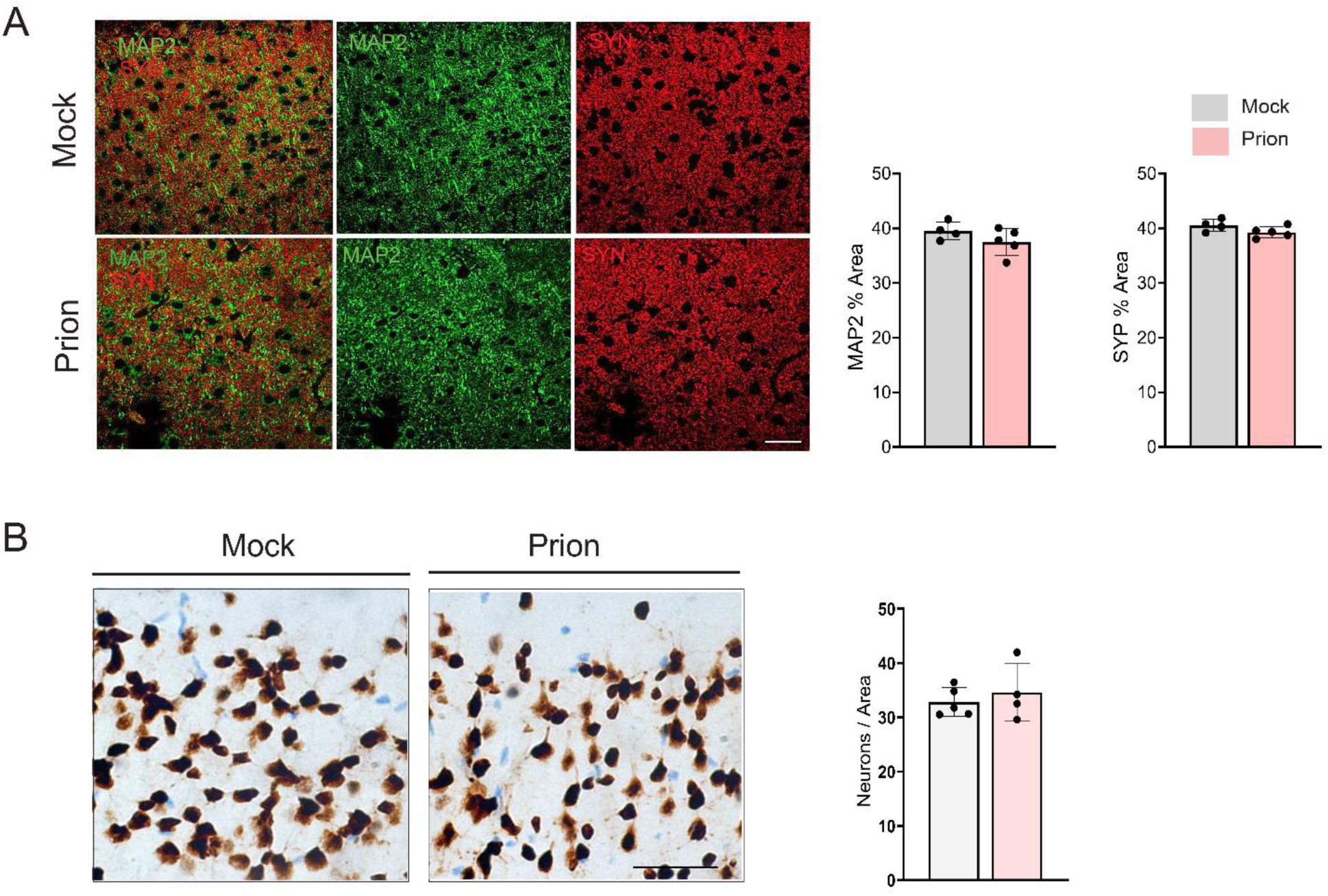
Dendrites, synapses, and neuronal numbers are unchanged in the prion-infected cortex at the 40% timepoint. (A) Representative confocal images of cortex immunolabeled for dendrites [MAP2 (green)] and pre-synapses [synaptophysin (red)]. Quantification (right) shows the percent area of fluorescent signal in cortical layers II-III. N = 4 (mock) and 5 (prion). Scale bar = 20 µm. (B) Representative images and quantification of neurons in cortical layers II-III immunolabeled for NeuN. N = 5 (mock) and 4 (prion), Scale bar = 100 µm. Bar graphs show mean ± SEM. Unpaired, 2-tailed *t* test with Welch’s correction.

**Figure 5-1.**
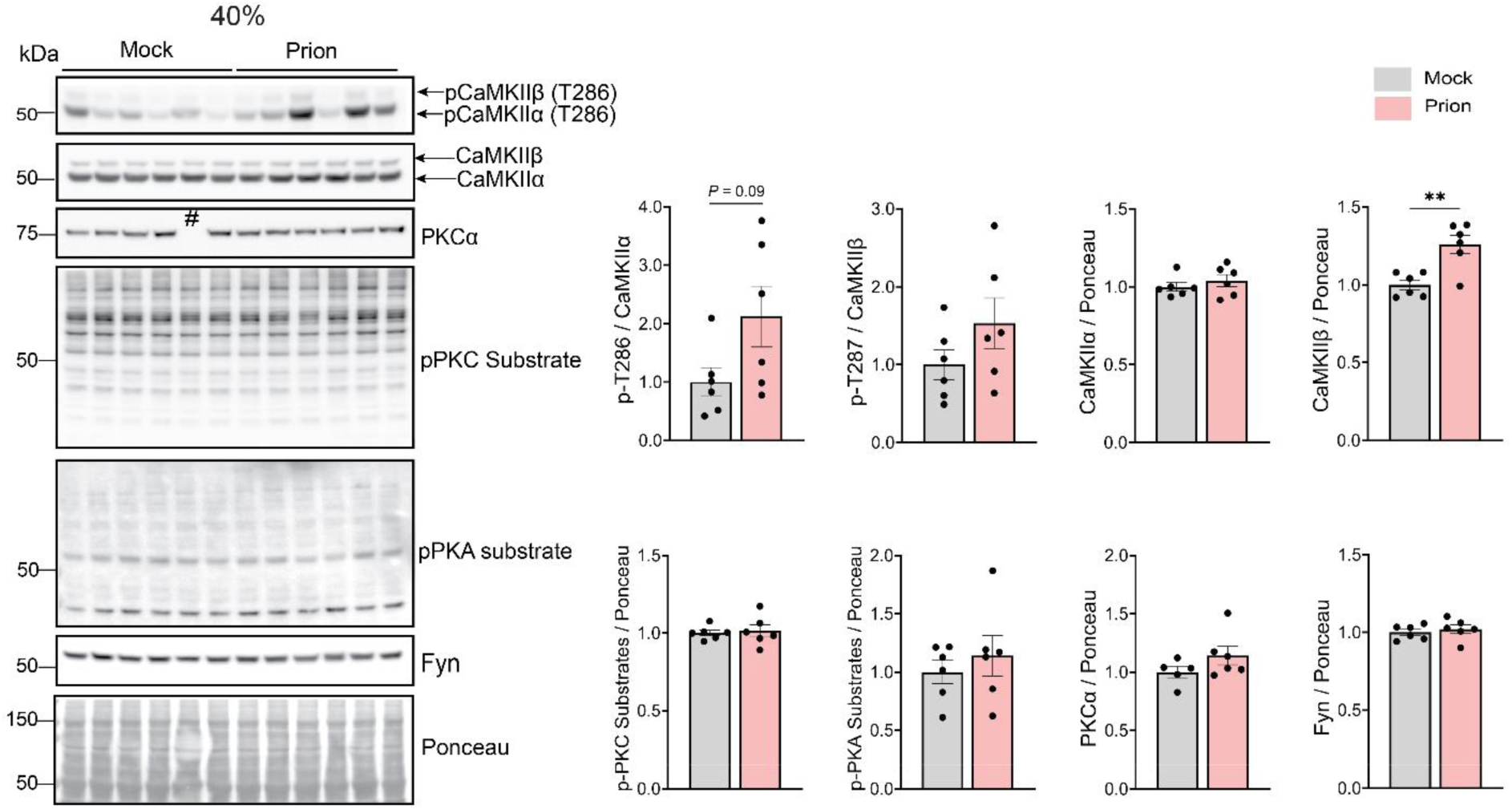
Kinase alterations in cerebral cortex at the 40% tp. Immunoblots and quantification of CaMKIIα/β, phospho-CaMKIIα/β-T286, PKC and PKA phosphorylated substrates, and total PKCα and Fyn levels. Mean ± SEM, N = 6 per group. Unpaired, 2-tailed *t* test with Welch’s correction. \*\**P*<0.01. For PKCα, one mock sample was not loaded (#). Ponceau corresponds to the pCaMKIIα/β (T286) membrane.

**Figure 4-1.**
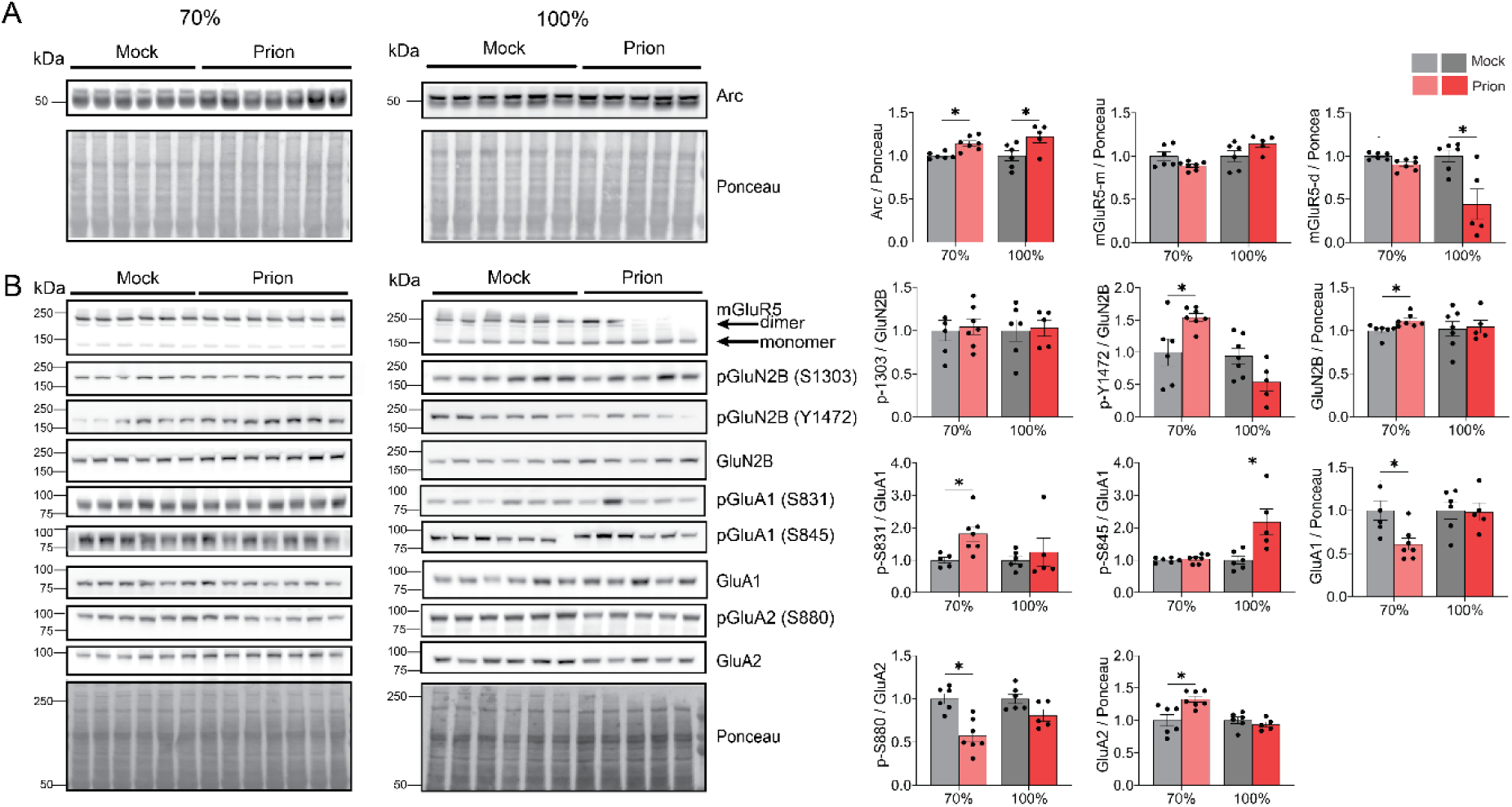
Phosphorylated GluA1 and GluN2B are increased at the clinical stages of prion disease. (A) Immunoblots and quantification of Arc at 70% and 100% relative to their respective mock controls. (B) Immunoblots and quantification of mGluR5 monomer and dimer, GluN2B and key phosphosites (S1303, Y1472, and S1480), and GluA1 and GluA2 and AMPAR phosphosites (GluA1: S845 and S831; GluA2: S880) at 70% and 100%. 70%: N = 6 (mock) and N = 7 (prion), 100% (terminal): N = 6 (mock) and N = 5 (prion). Mean ± SEM. Unpaired, 2-tailed *t* test with Welch’s correction. \**P*<0.05.

**Figure 6-1.**
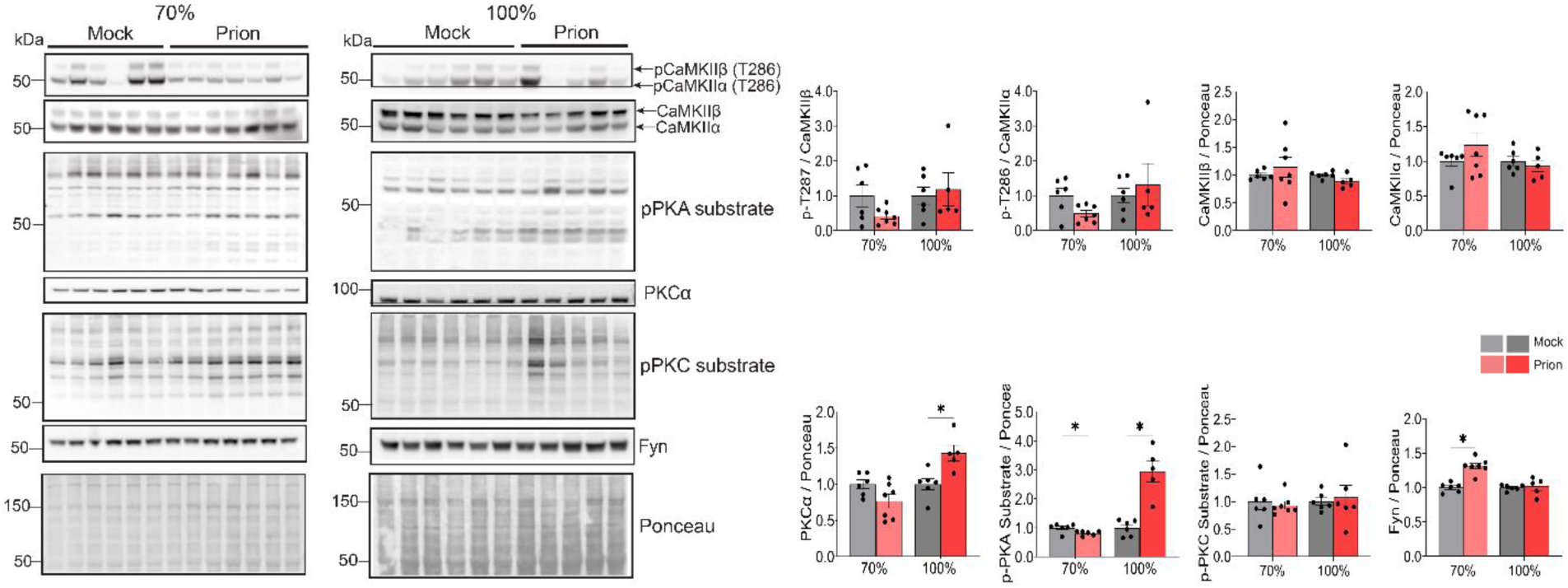
Kinase alterations at the clinical stages of prion disease. Immunoblots of CaMKIIα/β, phospho-CaMKIIα/β-T286, PKC and PKA phosphorylated substrates, and total PKCα and Fyn levels at 70% and 100% (left) and quantification (right). 70%: N = 6 (mock) and N = 7 (prion), 100% (terminal): N = 6 (mock) and N = 5 (prion). Mean ± SEM. Unpaired, 2-tailed *t* test with Welch’s correction. \**P*<0.05.

## Notes

### Competing Interest Statement

The authors have declared no competing interest.

### Funding

- National Institute of Neurological Disorders and Stroke: NS069566, NS076896, NS2033955, NS105498
- National Institute on Aging: 5T32AG066596
- University of California Office of the President: UC PresidentM-BM-^Rs Postdoctoral Fellowship Program
- CJD Foundation: The Harvey L. Hall Memorial Grant

